# Design of smart antibody mimetics with photosensitive switches

**DOI:** 10.1101/2020.12.03.410936

**Authors:** Lian He, Peng Tan, Yun Huang, Yubin Zhou

## Abstract

As two prominent examples of intracellular single-domain antibodies or antibody mimetics derived from synthetic protein scaffolds, monobodies and nanobodies are gaining wide applications in cell biology, structural biology, synthetic immunology, and theranostics. We introduce herein a generally-applicable method to engineer light-controllable monobodies and nanobodies, designated as moonbody and sunbody, respectively. These engineered antibody-like modular domains enable rapid and reversible antibody-antigen recognition by utilizing light. By paralleled insertion of two LOV2 modules into a single sunbody and the use of bivalent sunbodies, we substantially enhance the range of dynamic changes of photo-switchable sunbodies. Furthermore, we demonstrate the use of moonbodies or sunbodies to precisely control protein degradation, gene transcription, and base editing by harnessing the power of light.

## 1. INTRODUCTION

Intracellular single-domain antibodies (intrabodies) and their mimetics derived from synthetic protein scaffolds, as most notably exemplified by nanobodies and monobodies, are gaining wide use in cell biology, structural biology, synthetic immunology, and theranostics.^[1–6]^ Single-domain intrabodies and their mimetics such as monobodies or nanobodies rival conventional antibodies by their substantially smaller sizes (12-15 kDa vs 150-160 kDa), freedom from disulfide-bond formation, and ease of in vitro and in cellular expression. Recent engineering efforts have led to the generation of several classes of chemically or light-dependent single-domain antibodies, either based on split nanobodies (optobody)^[7]^ or hybrid proteins that utilize a photosensitive switch (OptoNB and OptoMB)^[8–9]^ or circularly permuted bacterial dihydrofolate reductase (LAMA).^[10]^ Nontheless, these engineered intrabodies exhibit a relatively slow activation kinetics or suffer from a relatively low or modest dynamic range of light-induced changes.

Bearing these unmet needs in mind, we introduce herein a set of smart monobodies and nanobodies that respond to light within seconds, in the form of ON-switches (sunbody) or OFF-switches (moonbody). The light-oxygen-voltage domain 2 (LOV2) derived from *Avena Sativa* phototropin 1 has been engineered into intrabodies to confer allosteric control of protein activity by light. We greatly improved the perfomance of our light-controlled moonbodies by optimzing the LOV2 insertion in the EF loop rather than the DE loop as described in the OptoMB tool^[9]^. By optimizing LOV2 insertion sites and paralleled insertion of two LOV2 modules into a single sunbody, we significantly enhanced the dynamic range of light-inducible response when compared to the existing OptoNB^[8]^. With these smart sunbodies or moonbodies, we demonstrate their wide applications in the remote control of protein localization, cell death, transcriptional programming and precise base editing.

## 2. RESULTS AND DISCUSSION

### 2.1 Design of Moonbody as Light-Controllable Antibody Mimetics

We first set out to design a light-controllable monobody (termed moonbody) by inserting the LOV2 photoswitch into a fibronectin type III domain (FN3) scaffold that specifically recognizes the Src Homology 2 (SH2) domain of Abelson tyrosine kinase (Abl).^[11–12]^ Six insertion sites at exposed loop regions (**Supplementary Figure 1a**) were selected, with three situated in the antigen-recognizing BC, DE, FG loops (the equivalent of complementarity-determining regions (CDRs) seen in an antibody) and the other three at the opposing loops (**Figure 1a-c**). We envisioned that light-induced conformational changes in LOV2 would allosterically modulate the moonbody-target interaction, thereby permitting photoswitchable control of the moonbody. To visualize the light-dependent changes in moonbody-SH2 association in live cells, we anchored the SH2 domain to the nuclear envelope (NE) via fusion with mEmerald-lamin A, and co-expressed the engineered moonbody as a cytosolic protein with partial distribution in the nucleoplasm (NP). Insertion of LOV2 at Site 4 (DE loop) abolished the moonbody-target interaction regardless of the presence of light (**Supplementary Figure 1b-c**), likely owing to the disruption of the antigen-binding pocket. The insertion of LOV2 at Site 2 (BC loop) led to an appreciable increase (<10%) of nucleoplasmic moonbody toward NE-SH2 (**Figure 1d**, and **Supplementary Figure 1b-c**). Contrariwise, insertion at flexible loop regions opposing the CDR-like areas, such as S1, S3, and S5, caused varying degrees of light-induced dissociation of moonbody from NE, with the S5 construct showing the highest dynamic change and hence used for further optimization (**Figure 1d**, and **Supplementary Figure 1a-c**).

**Figure 1.**
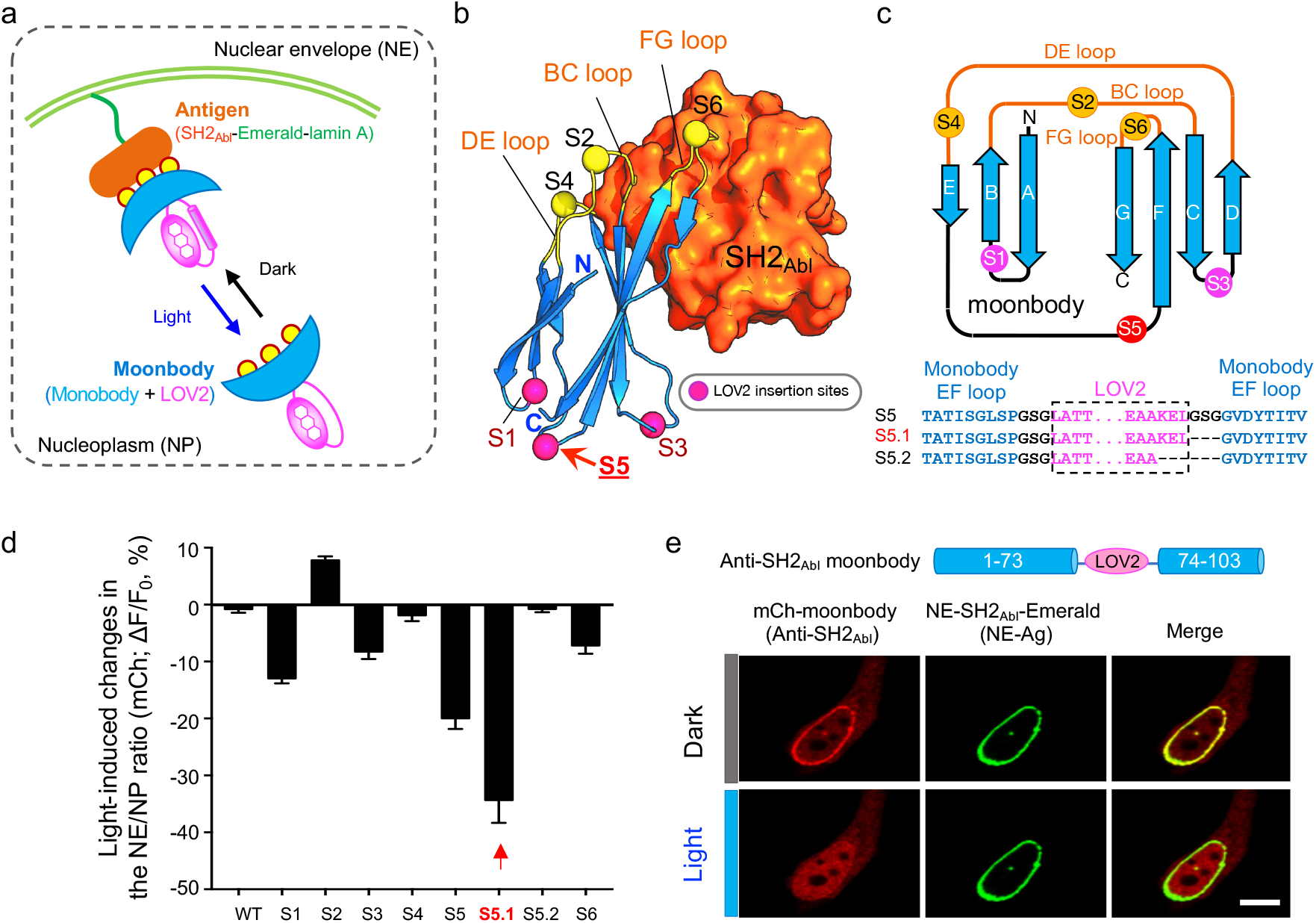
Design and optimization of light-switchable monobody (moonbody). a. Schematic depicting the design of light-switchable monobody (designated “moonbody”) and the nuclear envelope (NE) translocation assay used for screening. A photoswitch LOV2 is inserted into selected loop regions to enable photo-inducible target recognition in a reversible manner. Light-dependent shuttling of moonbody between NE and the nucleoplasm (quantified as the NE/NP ratio of mCh signals) is monitored. Yellow circles represent the three CDR (complementarity-determining region)-like loop regions that mediate moonbody-target recognition. b. LOV2 insertion sites mapped to the 3D structure of an anti-SH2Abi monobody (PDB entry: 3T04). Even-numbered insertion sites were created in the target-recognition loops, whereas odd-numbered sites were located opposite to the antigen-recognizing BC/DE/FG loops. c. 2D topology representation of an anti-SH2 monobody, with the insertion sites indicated by circles. The monobody-LOV2 junction regions for S5 or its variants were shown below the cartoon. See Supplementary Figure 1 for detailed sequence information of all constructs tested in the study. d. Quantification of light-dependent responses (as the NE/NP ratio) of moonbody variants. See Supplementary Figure 1 for representative images. Insertion at Site 5 (S5) led to the highest light-induced change. n = 6-25 cells. Data are shown as mean ± s.e.m.. e. Representative confocal images of a HeLa cell co-expressing an anti-SH2 moonbody (mCh-tagged variant S5.1; red) and NE-tethered SH2 domain of Abl kinase (NE tethered-antigen or abbreviated as NE-Ag; green) in the dark or after light illumination for 10 sec. Scale bar, 10 μm.

To further enhance the dynamic range of light-induced changes, we continued to optimize the linker regions flanking the insertion site and the C-terminal Jα helix within LOV2 (**Supplementary Figure 1a**), and ultimately identified one construct (S5.1; **Figure 1c**) showing almost complete dissociation from NE upon light stimulation (**Figure 1e**). We named this construct as “moonbody” because it retained its antigen-recognition ability in the dark, but rapidly dissociated from the antigen upon light stimulation. We further validated the transferability of our engineering approach by using two additional monobodies that recognize the small ubiquitin-like modifier protein (SUMO) and the maltose-binding protein (MBP).^[13–14]^. Both engineered moonbodies showed light-induced dissociation from their corresponding target proteins (**Supplementary Figure 2**). These moonbodies might find broad applications in the cost-effective elution or enrichment of recombinant MBP- or SUMO-fusion proteins by switching on and off light, respectively.

### 2.2. Spatiotemporal control of antigen recognition and target degradation with moonbody

The generation of moonbody allowed us to control antibody-antigen recognition with high spatial and temporal precision. We confirmed the feasibility of spatial control by alternatively focusing the photostimulation upon two individual cells under the same imaging field (**Figure 2a**). As anticipated, only the cell within the illuminated area showed light-dependent dispersion of NE-bound moonbody into the nucleoplasm; whereas the other cell without photostimulation showed no appreciable changes in the subcellular distribution of moonbody (**Figure 2a** and **Supplementary Movie 1**). When the whole imaging field was exposed to light stimulation, both cells showed simultaneous light-dependent moonbody redistribution (**Supplementary Movie 1**). Temporally, the moonbody-target interaction turned out to be reversible, as reflected by repeated NE-to-NP translocation of moonbody in response to at least 10 dark-light cycles of photostimulation (**Figure 2b** and **Supplementary Movie 2**). The activation and deactivation half-lives were determined to be 7.8 ± 0.1 s and 46.5 ± 0.3 s, respectively (**Supplementary Figure 1d**). Together, modular insertion of LOV2 into an FN3-derived moonbody enables noninvasive and reversible control of single-domain antibody mimetics by light.

**Figure 2.**
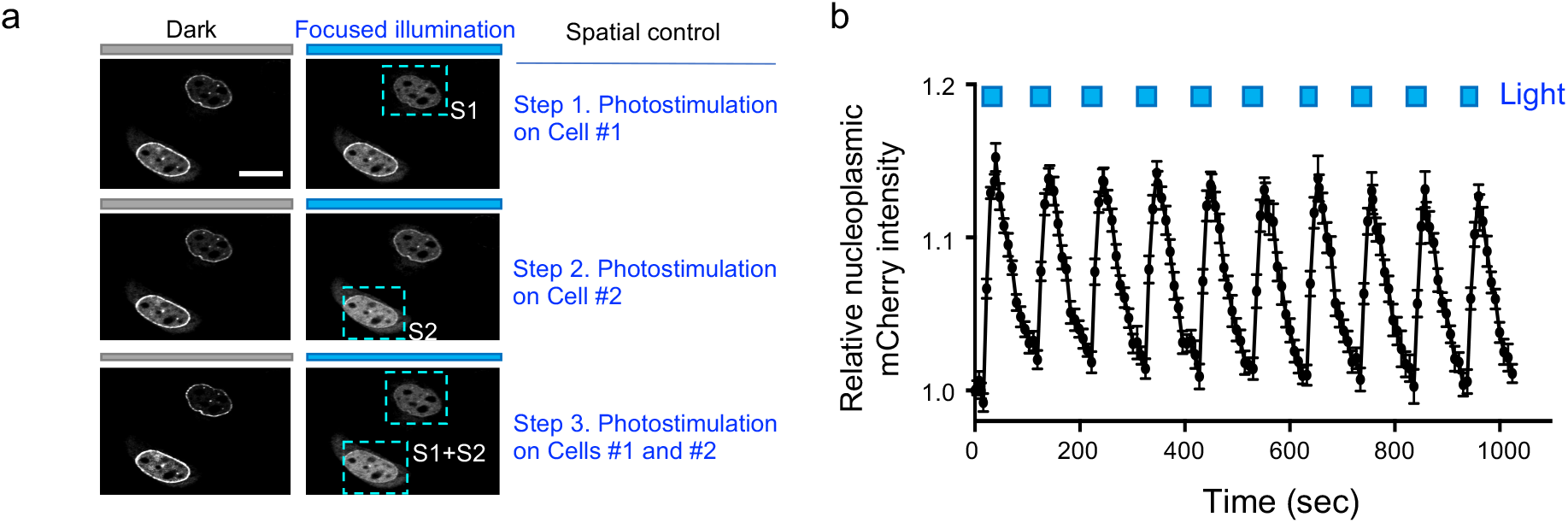
Spatiotemporal control of moonbody by light. a. Spatial control of the moonbody-antigen interaction in live cells. HeLa cells were cotransfected with NE-SH2_Abl_ (as the Ag; not shown) and mCh-moonbody (shown in gray). Photostimulation was sequentially applied to Cells 1 and 2 in the same imaging field as indicated by the blue box. Scale bar, 10 μm. Also see **Supplementary Movie 1**. b. Temporal control of the moonbody-antigen binding in live cells. The nucleoplasmic mCh intensity (as illustrated in panel b) in response to 10 repeated dark-light cycles of stimulation was quantified. Photostimulation was applied by using the 488-nm laser with 5% input. n = 11 cells. Also see **Supplementary Movie 2**.

We further asked whether moonbody could be utilized to conditionally fine-tune the expression levels of its binding target. To test this, we fused the SH2-specific moonbody with the auxin signaling F-box 2 protein (AFB2), a component in the Skp1-Cul-F-Box (SCF) E3 ubiquitin ligase complex that can recruit auxin-inducible degrons for proteasomal degradation^[15^] Moonbody can directly recognize its target, thus obviating the need for auxin and the fusion of the degron tag to a target protein. We reasoned that the light-triggered moonbody-target dissociation can prevent SH2 from being destroyed by the proteasomal degradation machinery (**Figure 3a**). Indeed, in the presence of escalating doses of pulsed light stimulation, we observed a gradual recovery of SH2-mEmerald signals in the transfected cells (**Figure 3b**), thus establishing the feasibility of using a photoswitchable moonbody to modulate the target protein expression levels in live cells.

**Figure 3.**
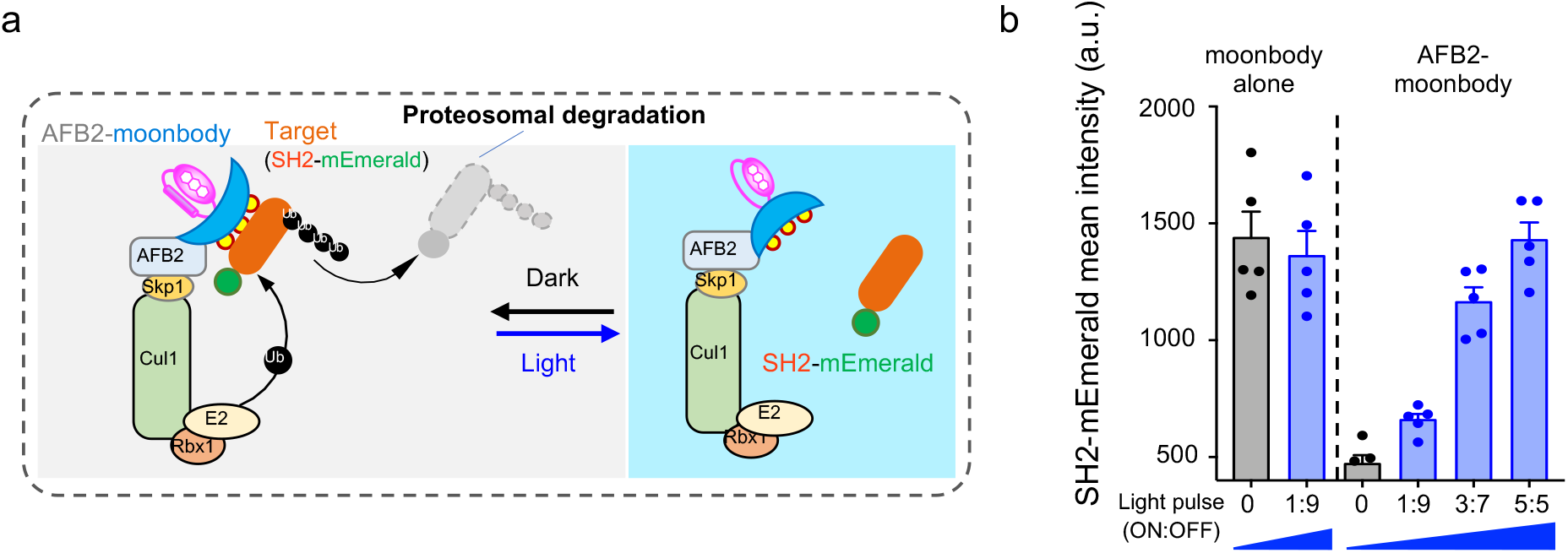
Light-tunable control of protein turnover by moonbody. a. Schematic illustrating the use of an anti-SH2 moonbody for light-tunable degradation of the target protein in mammalian cells. AFB2 binds the Skp1-Cul1-Rbx1 to form a ubiquitin ligase complex to mediate proteasomal degradation. Light-induced dissociation between the moonbody and its target can be exploited to conditionally control protein degradation. b. Quantification of light-tunable degradation of SH2-mEmerald using moonbody. HEK293 cells were transfected with AFB2-moonbody (or moonbody alone as control) and SH2-mEmerald, and then either shielded (Dark) or exposed to 8-h blue light illumination with intensifying pulses (with the ON and OFF durations indicated in the x-axis). An external 470-nm LED light was used as the light source (40 μW/mm^2^). n = 5 fields of view per condition. Error bars denote s.e.m..

### 2.3. Design of Sunbody as Photoactivatable Nanobody

We next extended the similar engineering approach, as well as the NE translocation assay, to screen photo-switchable nanobodies (designated “sunbodies”; **Figure 4a-b**). We used a mCherry-specific nanobody (LaM8)^[16]^ as a test case and inserted LOV2 into flexible loop regions opposing the CDRs (**Figure 4b** and **Supplementary Figure 3a**), the latter of which are involved in direct antigen recognition and thus remain unperturbed. The constructs S1, S2, and S4 exhibited negligible or little changes (<4%) in the NE translocation assay (**Supplementary Figure 3b-c**), making them less ideal for further optimization. We then resorted to two complementary approaches to enhance the light-dependent dynamic changes. First, because the N-terminus of nanobody is in close proximity to the CDRs (5-20 Å; S0 site; **Figure 4b**), we reasoned that direct fusion of LOV2 to the N-terminus of nanobody might partially cage its antigen-recognizing ability. Second, we expected that by insertion of LOV2 into the loop region (a potential allosteric site) positioned nearby CDR1 and CDR2 (S3 site; **Figure 4b**). These engineering efforts led to 10% and 22% increase in the NE/NP ratio after photostimulation, respectively (**Figure 4c** and **Supplementary Figure 3b-c**). These results further motivated us to test whether the simultaneous insertion of LOV2 at S0 and S3 could exert a synergistic effect on sunbody. Indeed, the resultant construct (S0 + S3) displayed over 80% changes in the NE/NP ratio (**Figure 4c**), and was designated as “sunbody” since its function can only be switched on in the lit condition. The engineered sunbody showed no appreciable accumulation at NE in the dark but exhibited strong light-triggered translocation toward NE (*t1/2*, on = 2.6 ± 1.8 s; **Figure 5**). The sunbody-antigen association was also found to be reversible (**Figure 5a**). Notably, the use of a dimeric concatemer of sunbody (2xsunbody) could further enhance the strength of light-switchable antibody-antigen binding, as reflected by >2-fold increase in the signal-to-background ratio reported by four additional subcellular translocation assays (from the cytosol to mitochondria [Mito], plasma membrane [PM], endoplasmic reticulum [ER], or early endosome [EE]; **Figure 5b-d** and **Supplementary Movies 3-4**). These findings clearly establish the feasibility of using sunbody to interrogate proteins located at different subcellular organelles.

**Figure 4.**
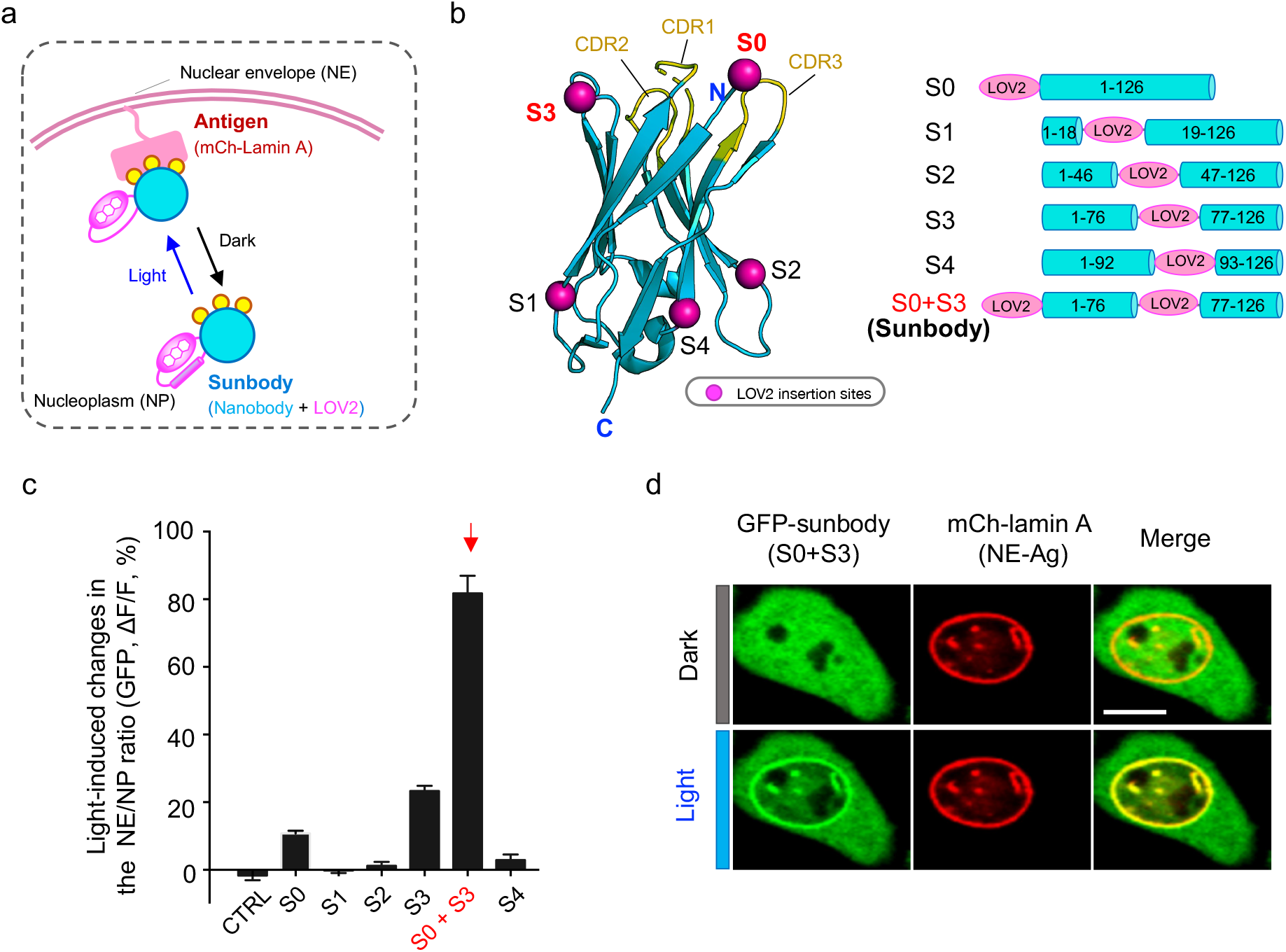
Engineering photoswitchable nanobody (sunbody) to enable light-controllable antigen binding. a. Cartoon depiction of the design and the NP-to-NE translocation assay. Photoswitchable redistribution of an engineered anti-mCherry (mCh) nanobody (designated “sunbody”) is used as the readout. Sunbody is expected to shuttle between NE and NP in a light-dependent manner. Yellow circles represent three CDRs involved in antigen binding. b. Insertion sites for LOV2 mapped to the modeled 3D structure of an anti-mCh nanobody (LaM8). S1, S2 and S4 are located at the opposite side of CDR loops. Both the N-terminus (S0) and S3 are in close proximity to CDRs. See Supplementary Figure 3 for detailed sequence information. c. Quantification of light-induced changes in the NE/NP ratio for an anti-mCh GFP-tagged sunbody. The combination of LOV2 fusion to the N-terminus (S0) and its additional insertion at S3 led to the strongest light-inducible changes (S0 + S3). See Supplementary Figure 3 for light-induced changes of each construct. n = 15-66 cells from three independent assays. Data are shown as mean ± s.e.m.. d. Representative confocal images of a HeLa cell co-expressing sunbody (GFP-tagged LaM8-S3; green) and NE-tethered mCh-lamin A (red) before and after light illumination for 10 sec. Scale bar, 10 μm.

**Figure 5.**
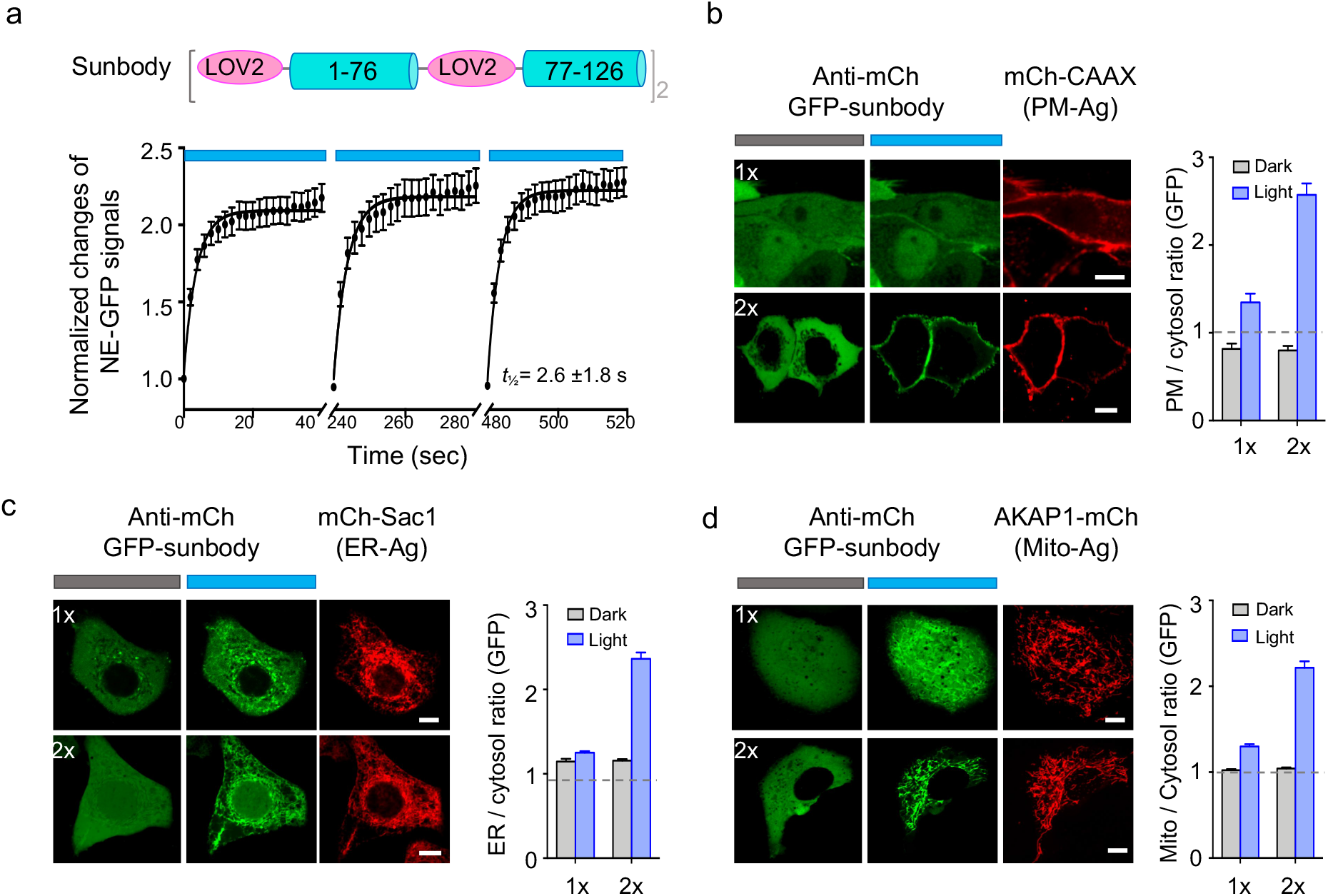
Improved sunbody shows high sensitivity for light-dependent subcellular targeting. a. Quantification of the sunbody-antigen interaction in response to three repeated dark-light cycles. The changes in the nucleoplasmic GFP signals were used as the readout. n = 23 cells. b-d. Sunbody used for light-dependent subcellular targeting of its binding partner. HeLa cells were transfected with an anti-mCh GFP-tagged sunbody (1x; green; top panels), or its concatemeric form (2x; green; bottom panels), along with the mCh as antigen (red) tethered to PM (b), ER (c), or outer mitochondrial membrane (d). The quantification of relative GFP signals at the corresponding subcellular organelles before and after light illumination were shown next to the images (n = 15-75 cells). The use of 2xsunbody in a single construct substantially enhanced the signal-to-noise ratio. Scale bar, 10 μm. Also see Supplementary Movies 3-4.

### 2.4. Sunbody for photo-controllable gene transcription and base editing

We then moved on to explore the use of sunbody for remote control of gene expression. We combined sunbody with the FLARE platform^[17]^ to create a SolarFLARE system for light-inducible transcriptional activation (**Figure 6a**). In our envisioned design, light stimulation initiates the translocation of a cytosolic anti-mCh sunbody-TEV hybrid protein toward PM-tethered mCh-FLARE components (mCh-LOV2-TCS-tetR-VP16), and brings TEV in close proximity to the exposed substrate (TEV cleavage site or TCS) to cleave the polypeptide, ultimately releasing the otherwise PM-restricted transcriptional coactivator (tetR-VP16) into the nuclei to recognize the nucleotide tetracycline operator (tetO) sequence and activate gene expression. The photosensitive LOV2 modules embodied in both sunbody and the FLARE system confer tight control over gene transcription by light. The photo-inducible gene transcription was first validated by using blue fluorescent protein (TagBFP) as a reporter (**Figure 6b**). In the dark, we did not observe appreciable TagBFP signals, attesting to the strict control of the SolarFLARE system. By contrast, a marked increase of TagBFP signals was noted in the light-illuminated group, suggesting that the light-inducible antibody-antigen interaction effectively activates the SolarFLARE system to drive gene expression (**Figure 6b**). We next replaced the TagBFP reporter with an N-terminal fragment of mixed lineage kinase domain-like pseudokinase (MLKL-NT; residues 1-182) that is capable of eliciting necroptotic cell death (necroptosis),^[18]^ with the goal of developing an optogenetic suicide device. Upon light stimulation, we observed a substantial increase of cell death, as reflected by the appearance of SYTOX blue staining of the nuclei of dead cells after PM permeabilization by MLKL-NT (**Figure 6c** and **Supplementary Figure 4**). These results establish SolarFLARE as a light-controlled transcriptional programming device.

**Figure 6.**
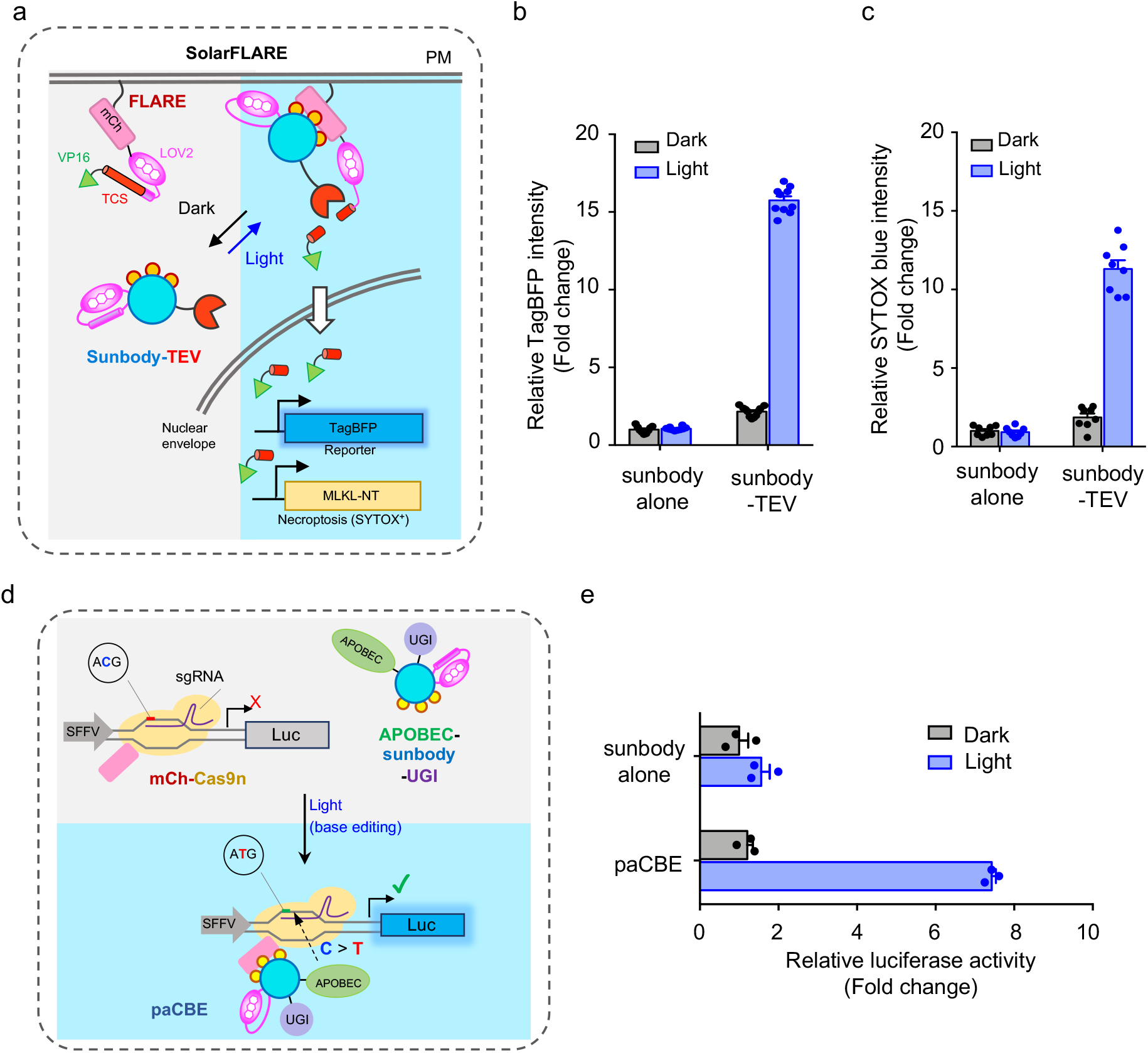
Use of sunbody to enable photoactivatable gene transcription and base editing. a. Cartoon illustrating the combination of sunbody with a modified FLARE system (designated SolarFLARE) to enable light-inducible expression of genes of interest, such as TagBFP as a reporter or the N-terminal domain of MLKL (MLKL-NT) as a necroptosis inducer. b. Quantification of BFP expression in HeLa cells transfected with SolarFLARE (sunbogy-TEV + FLARE) or the control (sunbody alone + FLARE) vectors, as well as the TagBFP reporter gene, before and after light illumination for 8 h. n = 10 fields of view from three independent assays. c. Quantification of necroptotic cell death as indicated by SYTOX blue nuclear staining of dead cells. HeLa cells were transfected with SolarFLARE (sunbogy-TEV + FLARE) or the control (sunbody alone + FLARE) vectors, as well as the inducible MLKL-NT expression cassette, before and after light illumination for 8 h. Also see Supplementary Figure 4 for representative images. n = 10 fields of view from three independent assays. d. Design of a photoactivatable cytosine base editor (paCBE). Upon photostimulation, sunbody-mCh association re-assembles two functional units of CBE (Part I: the mCh-Cas9n/sgRNA for genome targeting; Part II: APOBEC1-sunbody-UGI for C-to-T conversion) to restore the activity of paCBE. A “Gene ON” (GO) luciferase reporter system is used to report the activity of paCBE before and after light stimulation. Successful recruitment of Part II to the targeted genomic locus is anticipated to cause C-to-T conversion in the start codon (ACG>ATG) to initiate the translation of a luciferase reporter gene. e. Quantification of the base editing efficiency of paCBE by using luciferase activity as readout. Sunbody alone was used as a negative control. n = 3 independent assays.

Next, we set out to combine sunbody with the CRISPR/Cas9-mediated C-to-T base editing technique^[19]^ to design a photoactivatable cytosine base editor (paCBE). To achieve this, we split the functional units of CBE into two parts (**Figure 6d**): Part I contains a mCherry-tagged partially enzymatically disabled Cas9 (Cas9 nickase, or Cas9n) with sgRNA (mCh-Cas9n); whereas Part II contains anti-mCh sunbody fused with the cytidine deaminase APOBEC1 and a uracil DNA glycosylase inhibitor (UGI) to prevent U:G mismatch being repaired back to a C:G base pair^[20]^ We reasoned that, upon light-triggered sunbody-mCh interaction, paCBE could restore its cytosine-to-thymine editing function. To quickly test this idea, we used a “Gene ON” (GO) luciferase reporter system^[21]^ to monitor the activity of paCBE before and after light stimulation (**Figure 6e**). The GO system becomes activated only when a functional CBE converts C to T to create a *de novo* ATG codon at the beginning of a reporter gene (e.g., luciferase), thereby enabling translational initiation and successful production of the gene. When the transfected cells were shielded from blue light, we did not observe notable bioluminescent signals. Upon photostimulation, we detected a substantial increase of bioluminescence, presumably owing to the expression of luciferase after C>T conversion at the start codon (**Figure 6e**). Collectively, our results indicate the successful design of a photoswitchable base editing system by taking advantage of the light-inducible sunbody-antigen interaction.

## 3. CONCLUSIONS

In summary, by using widely-applicable intrabodies as protein scaffolds, we have illustrated the successful design of a series of synthetic sunbodies and moonbodies, in the form of ON-switches (sunbody) or OFF-switches (moonbody), that respond to light within dozens of seconds. Complementary to the recently developed optobodies, optoNBs and LAMAs,^[7–8, 10]^ our engineering efforts greatly expand the smart antibody toolbox by (i) extending the optogenetic engineering approach to antibody mimetics, and (ii) offering switchable intrabodies with varying kinetic and dynamic features. Worthy to note, we have enhanced the range of dynamic changes of photo-switchable sunbodies (over 2-fold compared to optoNB) by taking two approaches: paralleled insertion of two LOV2 modules into a single sunbody and the use of sunbody concatemers. The perfomance of moondies (optimized LOV2 insertion in the EF-loop) has been greatly improved compared to the recenlty reported OptoMB. Given the high modularity and facile transferability of our engineering approaches, we believe that a growing number of smart antibodies and antibody mimetics can likewise be generated in the near future. These molecular tools will greatly facilitate the mechanistic dissection of cell signaling and accelerate the development of personalized medicine, such as chimeric antigen receptor (CAR) T-cells and customized therapeutic biologics that respond to drugs and beverage intake. In particular, beverage-switchable antibodies are of immediate translational values given their high compatibility with in vivo applications and the relatively low barrier toward clinical trials.

## 4. EXPERIMENTAL SECTION

### Cell lines and culture conditions

HeLa and HEK293T (human embryonic kidney) cell lines were obtained from ATCC and cultured under 37 °C at a 5% CO_2_ atmosphere, and maintained in the Dulbecco’s modified Eagle’s media (DMEM, Sigma-Aldrich, St Louis, MO, USA), supplemented with 10% fetal bovine serum (FBS).

### Molecular cloning and plasmid construction

The standard restriction enzyme digestion-ligation and NEBuilder HiFi DNA Assembly methods were used to create plasmids in this study. The KOD Start DNA polymerase (EMD Millipore, MA, USA) was used for PCR amplification. All the subcloned sequences were verified using diagnostic restriction digestion and Sanger’s sequencing analysis. All the other cloning reagents were purchased from New England Biolabs (Ipswich, MA, USA).

cDNA sequences encoding monobodies and nanobodies used in this study were codon-optimized and synthesized as gBlock by Integrated DNA Technologies (IDT Inc, IA, USA). The monobody-encoding cDNAs (SH2_Abl_, MBP and SUMO) were individually inserted into a customized pcDNA-mCherry vector between the EcoRI and XbaI restriction sites to generate mCh-tagged monobodies. To create a nuclear envelope (NE)-targeting SH2_abl_, cDNA of lamin A was inserted into a customized mEmerald-C1 vector, followed by SH2_abl_ insertion (NheI-EcoRI). AsLOV2 fragments were PCR amplified and inserted into monobodies by using the NEBuilder HiFi DNA Assembly method. For photoswitchable degradation, moonbody (S5.1) cDNA was amplified via PCR and then inserted into the pSH-EFIRES-P-AtAFB2-mCherry vector (Addgene, #129716) between EcoRI and NotI sites to replace mCherry.

To make GFP fused nanobody, cDNA encoding the anti-mCherry nanobody LaM8 was cloned into the pTriEx-GFP vector between HindIII and XhoI sites to yield pTriEx-GFP-LaM8. AsLOV2 fragments were PCR amplified and inserted via NEBuilder HiFi DNA Assembly. The construct exhibiting the highest light-induced changes was designated as “sunbody” (S0+S3). The tandem sunbody expression vector (2xsunbody) was made by inserting one additional copy of sunbody into the pTriEx-GFP-sunbody plasmid. To make mitochondria-targeting mCh, the cDNA sequences encoding human AKAP1_1-30_ (flanked by NheI and BamHI) were inserted into mCherry-N1 to yield AKAP1-mCh. For ER anchored mCh, the human Sac1 fragment (residues 521-587) was cloned into the pEGFP-C3 backbone by utilizing the EcoRI-KpnI restriction sites, followed by GFP replacement by mCh (between NheI and XhoI sites). The plasma membrane targeting mCh construct was made as AgeI-mCh-EcoRV-CAAX-XbaI in the same backbone.

To create a SolarFLARE system for light-inducible transcriptional activation, the TEV protease component (from Ca-FLARE (protease), Addgene #92214) was assembled into pTriEx-GFP-sunbody to yield GFP-sunbody-TEV. The mCh and LOV2-TCS (TEV cleavage site)-tetR-VP16 fragments (from Ca-FLARE (TF), Addgene, # 92213) modules were fused into a pcDNA3.1(+) backbone to make PM-anchored mCh-LOV2-TCS-tetR-VP16. The TagBFP or MLKL expression cassette used in the SolarFLARE system was made by putting TagBFP cDNA or human MLKL-NT (1-182) (EcoRI/XbaI) under the control of a tight TRE promoter.

For photoactivatable cytosine base editor (paCBE), mCh and Cas9n (Cas9-D10A nickase) fragments were inserted into a pcDNA3.1(+) vector via HiFi assembly to make mCh-Cas9n (Part I). FNLSHiFi was replaced by GFP-sunbody in the pLenti-FNLSHiFi-P2A-Puro vector (Addgene, #136902) to make the APOBEC1-GFP-sunbody-UGI fusion construct (Part II). The luciferase-based GO system with sgRNA in the same vector was obtained from Addgene (pLenti-mU6-Luc2GO-PGK-Neo, Addgene #136905).

### Cell transfection

DNA transfection was performed by using the Lipofectamine 3000 transfection reagent (Thermo Fisher Scientific, MA, USA) according to the manufacturer’s instructions. For live-cell fluorescence imaging experiments, cells were seeded in four-chamber 35-mm glassbottom dishes (D35C4-20-1.5-N, Cellvis, Mountain View, CA, USA) one day before transfection, and imaged 24-48 h after transfection in an imaging cage equipped with 5% CO2 with the temperature set at 37 °C.

### Live cell photostimulation, time-lapse imaging and imaging data analysis

Time-lapse confocal imaging was performed on a Nikon A1R confocal module mounted onto an inverted Nikon Eclipse Ti-E body. The light sources came from a multi-line argon laser module containing 405, 488, 561 and 640 nm lasers. A live-cell imaging caged platform was used to maintain the temperature at 37 °C with 5% CO2 to keep cells healthy during the imaging process. A 10x, 20X air objective lens and 40x or 60x oil immersion objective lens were used for image acquisition.

To monitor light-induced association or dissociation between sunbodies/moonbodies and the corresponding antigens, HeLa cells seeded on glass-bottom dishes were co-transfected with the indicated plasmids shown in the related figures. Confocal images were acquired 24-48 hours after transfection. The cells were imaged every 4 sec for 2nmin unless otherwise noted. The 488-nm laser source to excite GFP was also used for photostimulation (with 1-5% output). To quantify fluorescent signals at selected areas, we used the region-of-interest (ROI) toolbox in Nikon NIS-Elements software to define the nuclear envelope (NE) or nucleoplasm (NP) regions. The “Time Measurement” tool was used to determine the mCherry intensities for moonbody variants and GFP intensities for sunbody variants. The fluorescence intensity ratio (F_NE_/F_NP_) was used as readout, with the changes in the ratio plotted as F/F_0_ or ΔF/F_0_. For spatially-restricted photostimulation, the FRAP module in the Nikon imaging system was used, with the 488-nm laser power output set at 0.2%-5%.

### Moonbody regulated protein degradation

The plasmid encoding the moonbody fused with the F-box protein atAFB2 was co-transfected with SH2-mEmerald into HeLa cells. Cells were treated with or without a customized blue light source (470 nm, 40 μW/mm^2^) after 16 h transfection. Light stimulation was applied for 10% (1 min ON, 9 min OFF), 30% (3 min ON, 7 min OFF), or 50% (5 min ON, 5 min OFF). After an additional 16 hours, cells were imaged and SH2-mEmerald mean intensity was measured. Moonbody without atAFB2 was used as control.

### SolarFLARE system for gene expression

The construct encoding GFP-sunbody fused with TEV protease (GFP-sunbody alone as a control) was co-transfected with PM-mCh-LOV2-TCS-tetR-VP16 and the pTRE-TagBFP reporter in HEK293T cells. Eight hours post-transfection, cells were exposed to pulsed blue light stimulation for 16 h (470 nm, 40 μW/mm^2^) with a light cycle of 1 min ON and 9 min OFF. Cells kept in the dark was used as a control. Confocal images were acquired after 48 hours of transfection with a 10x or 20x air object for mCh, GFP and BFP channels. Eight fields of view were recorded for each condition. GFP expression area was masked and the mean TagBFP intensity was calculated in this mask, with areas outside GFP-positive areas used as background. The background-corrected mean TagBFP intensities were calculated and plotted. For light-inducible necroptosis, the pTRE-TagBFP vector was replaced by the pTRE-MLKL-NT plasmid, with the remaining procedures and conditions identical to the SolarFLARE-BFP reporter experiment described above. To monitor cell death in real-time, live cells were stained with the SYTOX blue dye (Thermo Fisher Scientific, S11348, 1:5000 dilution, C_f_= 1 μM).

### Luciferase reporter assay

To examine the efficiency of photoactivatable cytosine base editor (paCBE), the mCh-Cas9n, APOBEC-sunbody-UGI, and pLenti-mU6-Luc2GO-PGK-Neo (as base editing reporter) were co-transfected into HEK293T cells. Cells were treated with blue light (470 nm, 40 μW/mm^2^, 1 min ON-9 min OFF cycles for 16 h) or kept in the dark 8 hours after transfection. Sunbody alone to replace APOBEC-sunbody-UGI was used as control. 72 h post-transfection, bioluminescence measurements were performed by using a Bright-Glo Luciferase Assay System from Promega (catalog #: E2610) by directly adding reagents to the culture medium at a 1:1 ratio. Five minutes later, the luminescence signals were quantified by using a Cytation 5 Cell Imaging Multi-Mode Reader (BioTek, Winooski, VT, USA).

### Statistical analysis

All the data were plotted using the GraphPad Prism 8.3.0 graphing software. Quantitative data were shown as mean ± s.e.m. unless otherwise noted. The analyzed number (n) of samples were described in the figure legends for each experiment. The half-lives and median effective concentrations (EC_50_) were determined by using the GraphPad Prism software package.

## ACKNOWLEDGEMENTS

We thank the financial supports from the National Institutes of Health (R01GM112003 to YZ, R21GM126532 to YZ, R01CA232017 to YZ, R01HL134780 to YH, and R01HL146852 to YH), the Welch Foundation (BE-1913-20190330 to YZ), the John S. Dunn Foundation (to YZ), and the American Cancer Society (RSG-16-215-01-TBE to YZ and RSG-18-043-01-LIB to YH).

## AUTHOR CONTRIBUTIONS

YZ and YH conceived the ideas and directed the work. LH, PT and YZ designed the study. LH and PT performed the experiments. LH and PT analyzed the results. YZ, YH and LH wrote the manuscript. All of the authors contributed to the discussion and editing of the manuscript.

## COMPETING INTERESTS

The authors declare no competing interests.

## Supplementary materials contain

Supplementary Figs. 1-4

Supplementary Movies 1-4

## Captions for Supplementary Movies

**Supplementary Movie 1. Spatial control of the moonbody-antigen interaction in mammalian cells.** Time-lapse imaging of two HeLa cells co-expressing anti-SH2_Abi_ mCherry-moonbody (shown in grey) and the nuclear envelope-tethered antigen (SH2_Abl_-mEmerald-lamin A; not shown here). Sequential localized photostimulation (488 nm laser; 0.5% output) was applied as follows: (1) Cell #1 at the top-right corner (0-12 s OFF; 16-49 s ON; 59-129 s OFF); (2) Cell #2 at the bottom-left corner (139-180 s ON; 182-262 s OFF); and (3) both cells (271-290 s ON; 300-390 s OFF).

**Supplementary Movie 2. Temporal control of the moonbody-antigen interaction in live cells.** Time-lapse imaging of two HeLa cells co-expressing an anti-SH2 mCherry-moonbody (grey) and the nuclear envelope-tethered antigen (SH2-mEmerald-lamin A; not shown). Pulsed photostimulation at 488 nm was applied as indicated by the vertical bars on the right graph. The fluorescent intensities of two circled regions in both cells were plotted on the right.

**Supplementary Movie 3. Time-lapse imaging of a HeLa cell co-expressing the antigen (Mito-mCh; red, middle) and an anti-mCherry, GFP-tagged 2xsunbody (green, right)**. A 488 nm-laser with 1% output was used for photostimulation.

**Supplementary Movie 4. Light-induced protein translocation to different subcellular organelles.** Shown was timelapse imaging of HeLa cells co-expressing an anti-mCherry, GFP-tagged 2xsunbody (green, middle panels) and mCherry as the antigen (red, right panels), tethered to ER (top panels), PM (middle panels) or early endosome (bottom panels). A 488 nm-laser with 5% output was used for photostimulation.

## Notes

### Competing Interest Statement

The authors have declared no competing interest.

