## Supplementary Figs for "Design of smart antibody mimetics with photosensitive switches"

**Contents:**

Supplementary Figures 1-4

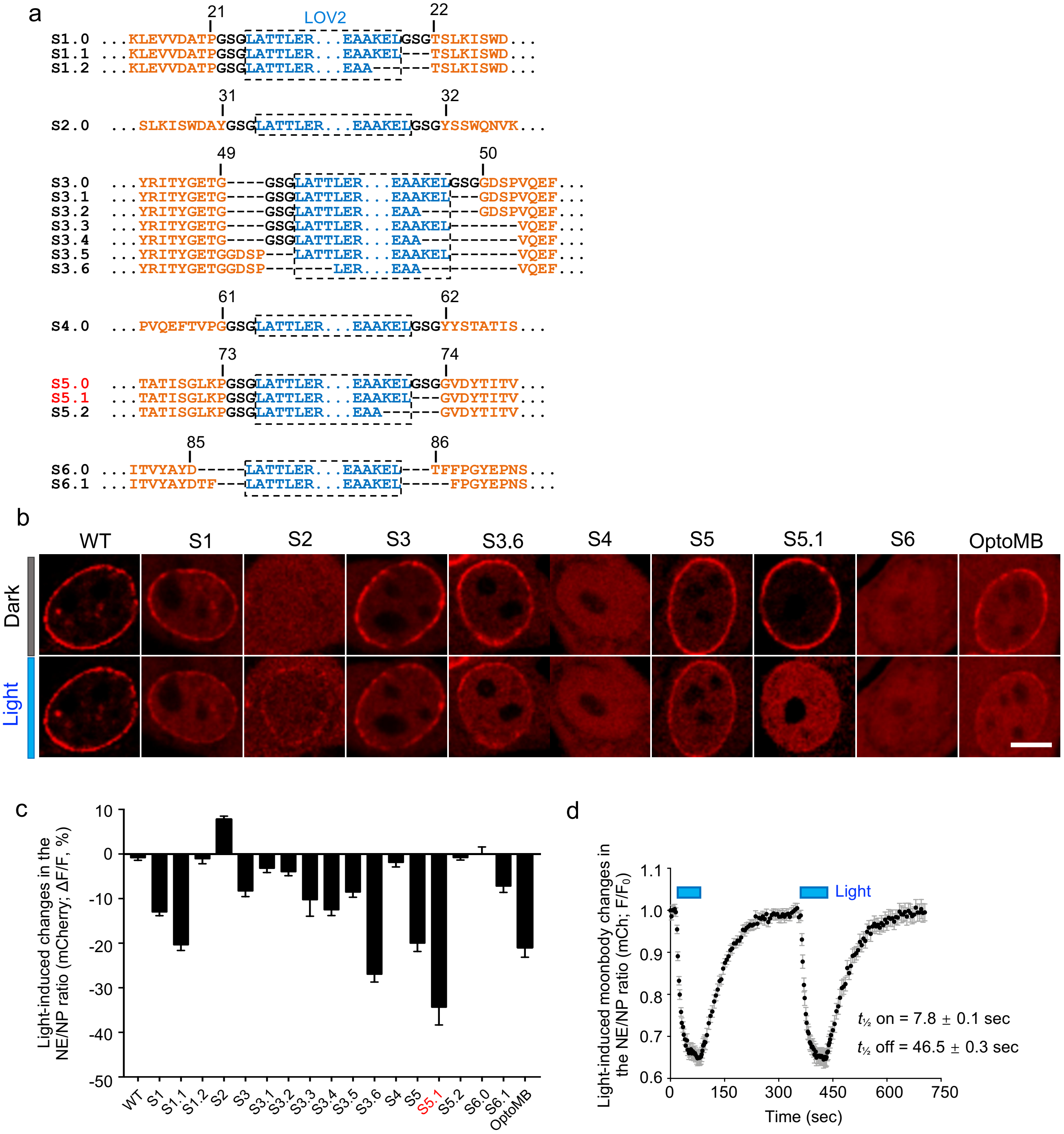


**Supplementary Figure 1 | The design and optimization of moonbody variants.**

a. Sequence information for moonbody variants tested in the study. The monobody-LOV2 junction regions were shown for each construct.

b. Representative confocal images of HeLa cells co-expressing the antigen (NE-SH2; not shown) and the indicated mCh-moonbody variants (red) before and after photostimulation at 470 nm for 120 sec. Scale bar, 10 µm.

c. Summary of the degrees of dissociation (quantified by the NE/NP ratio of mCh signals) from the target (NE-SH2) for the indicated moonbody variants. Selected data were also presented in **Fig. 1d**. n = 6-25 cells (mean ± sem).

d. Quantification of reversible changes in the NE/NP ratio. HeLa cells co-expressing the anti-SH2 moonbody and NE-tethered SH2 were subjected to two light-dark cycles of stimulation. n = 27 cells (mean ± sem).

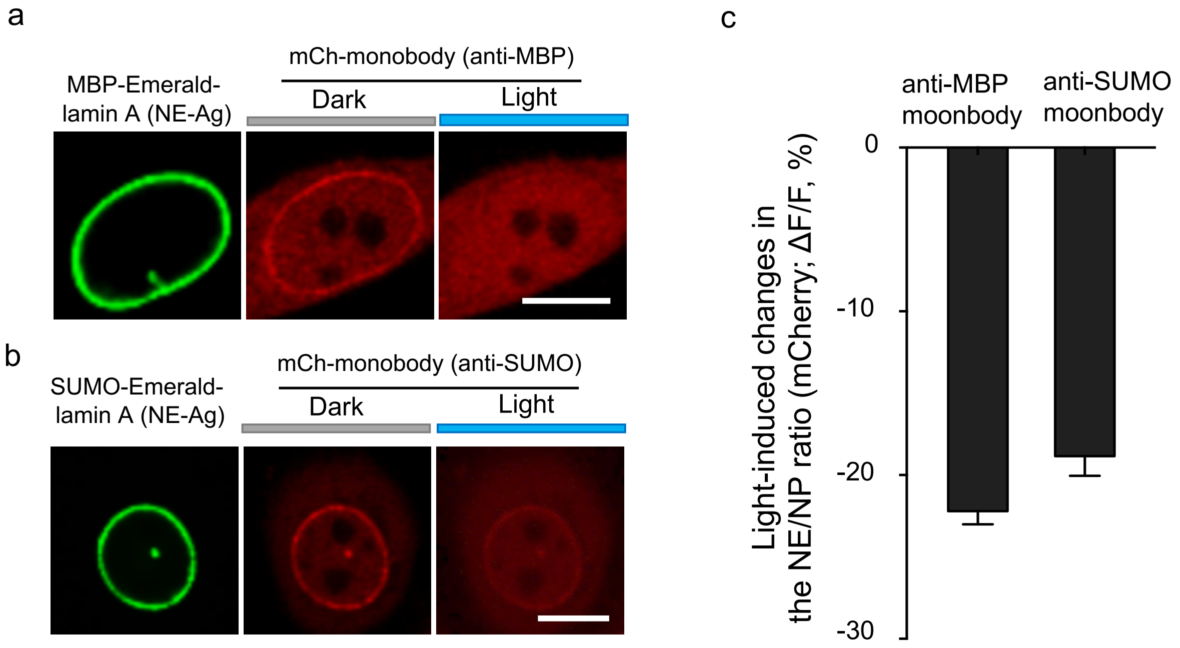


**Supplementary Figure 2 | Characterization of moonbodies against SUMO and MBP.**

a. Confocal images of a HeLa cell co-expressing an anti-MBP moonbody (mCh-tagged; red) and NE-tethered GFP-MBP in the dark or after blue light illumination for 120 sec at 470 nm. Scale bar, 10 µm.
b. Confocal images of a HeLa cell co-expressing an anti-SUMO moonbody (mCh-tagged; red) and NE-tethered GFP-SUMO (green) before (dark) and after blue light illumination for 120 sec at 470 nm. Scale bar, 10 µm.
c. Quantification of light-dependent response (as the NE/NP ratio of mCh signals) of the MBP or SUMO-specific moonbodies. n = 33-67 cells from three independent assays (mean ± sem).

a

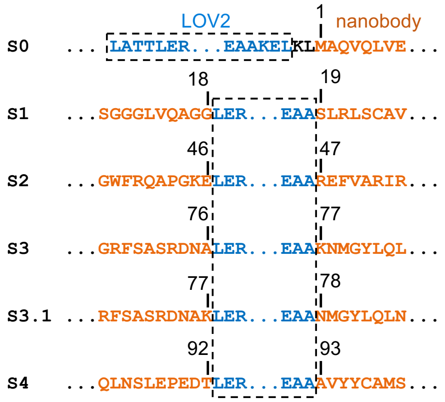


b

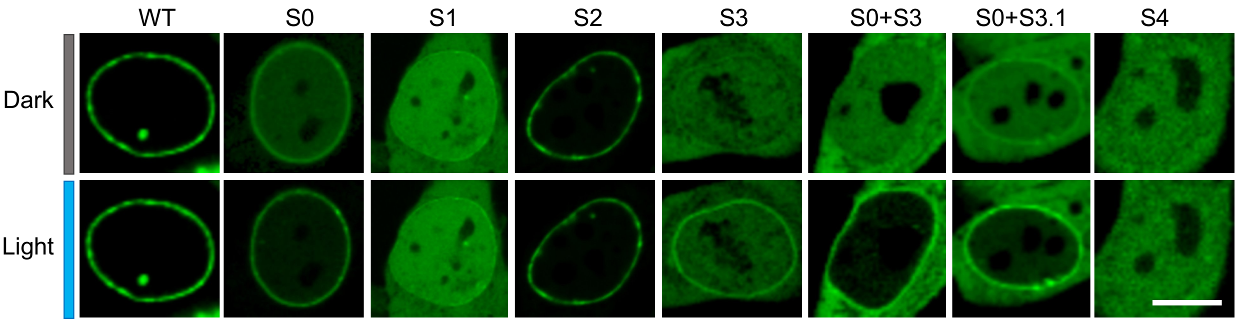


c

| Sunbody variants | Target binding (dark) | Light-induced change |
|---|---|---|
| WT | Strong | - |
| S0 | Strong | ++ |
| S1 | Weak | - |
| S2 | Strong | - |
| S3 | Weak | +++ |
| S0 +S3 | No binding | +++++ |
| S0 + S3.1 | Weak | ++++ |
| S4 | No binding | - |

**Supplementary Figure 3 | The design and optimization of sunbody variants.**

a. Sequence information of sunbody variants tested in the study. The sunbody-LOV2 junction regions were shown for each construct.

b. Representative confocal images of HeLa cells co-expressing the NE-tethered mCherry (as antigen; not shown) and the indicated GFP-sunbody variants (green). Scale bar, 10 µm.

c. Summary of the degrees of antigen (mCh) binding for the indicated GFP-sunbody variants before and after light stimulation (see panel **c**). -, <5%; +, 5-10%, ++, 10-20%; +++, 20%-40%; ++++, 40%-60%; +++++>60% increase in the NE/NP ratio.

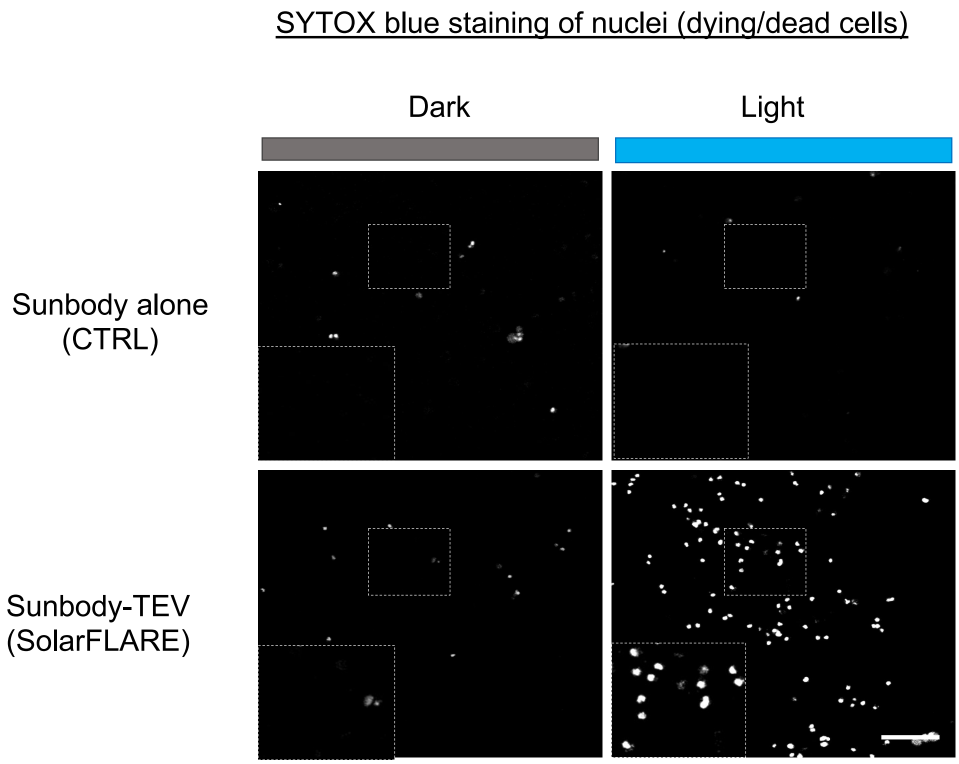


**Supplementary Figure 4 | A SolarFLARE system tailored for light-inducible MLKL-NT expression to induce necroptosis.** Representative confocal images of HeLa cells transfected with the SolarFLARE system (sunbody-TEV + FLARE; bottom), or the sunbody alone + FLARE combination as control (top), before and after light stimulation. SYTOX Blue staining was used to report dying cells. The quantification results were shown in **Fig. 6c**. Scale bar, 100 μm.
